# The effect of prior long-term recellularization with keratocytes of decellularized porcine corneas implanted in a rabbit anterior lamellar keratoplasty model

**DOI:** 10.1101/2020.12.31.424936

**Authors:** Julia Fernández-Pérez, Peter W. Madden, Robert Thomas Brady, Peter F. Nowlan, Mark Ahearne

## Abstract

Decellularized porcine corneal scaffolds are a potential alternative to human cornea for keratoplasty. Although clinical trials have reported promising results, there can be corneal haze or scar tissue. Here, we examined if recellularizing the scaffolds with human keratocytes would result in a better outcome. Scaffolds were prepared that retained little DNA (14.89 ± 5.56 ng/mg) and demonstrated a lack of cytotoxicity by *in vitro*. The scaffolds were recellularized using human corneal stromal cells and cultured for between 14 in serum-supplemented media followed by a further 14 days in either serum free or serum-supplemented media. All groups showed full-depth cell penetration after 14 days. When serum was present, staining for ALDH3A1 remained weak but after serum-free culture, staining was brighter and the keratocytes adopted a native dendritic morphology with an increase (p < 0.05) of keratocan, decorin, lumican and CD34 gene expression. A rabbit anterior lamellar keratoplasty model was used to compare implanting a 250 µm thick decellularized lenticule against one that had been recellularized with human stromal cells. In both groups, host rabbit epithelium covered the implants, but transparency was not restored after 3 months. Post-mortem histology showed under the epithelium, a less-compact collagen layer, which appeared to be a regenerating zone with some α-SMA staining, indicating fibrotic cells. In the posterior scaffold, ALDH1A1 staining was present in all the acellular scaffold, but in only one of the recellularized lenticules. We conclude that recellularization with keratocytes alone may not be sufficiently beneficial to justify introducing allogeneic cells without concurrent treatment to further manage keratocyte phenotype.

## 1 Introduction

The limited supply of human corneas for transplantation requires investigation of an alternative supply and one promising solution has been the use of decellularized corneas [1] [2] [3] [4]. These corneas may be obtained from human donors whose corneas would otherwise be unsuitable for transplantation [5], but an animal source such as the pig offers a much greater supply of tissue. Decellularized porcine corneal stroma provides a collagen-proteoglycan scaffold with appropriate composition and architecture that inherently supports corneal transparency and cellular repopulation. These corneas have already undergone clinical trials with encouraging results [1] [2] [5] [3] [4]. The laboratory work that preceded these trials included implantation of both human or porcine decellularized corneal stroma, primarily using a rabbit model of both interlamellar pocket and anterior keratoplasty [6] [7] [8] [9] [10] [11]. These studies indicated that although neovascularization and immune reaction was slight, corneal haze did hamper vision for some time, although corneal transparency eventually began to return as the study length increased [3]. These findings support the concept that decellularized corneal scaffold serves as a template for remodelling [6], but there is always a delay in transparency, or the risk of permanent haze from scar tissue. Although the epithelium rapidly returned to the anterior surface of the scaffold, few keratocytes repopulated the stroma, even after 12 months [7] [8] [12] [9]. Such a long delay appears undesirable, as one role of keratocytes is to remodel the stromal extracellular material (ECM) and thereby regulate collagen fibril spacing to restore corneal transparency [13]. However, it has been recognised that even disordered collagen fibrils in ECM can still be transparent [14] if they are not too thick, which is the case with porcine derived scaffold.

Clinically, if the cornea is damaged leaving the ECM intact, quiescent corneal keratocytes proliferate without phenotype switching or scar formation. [15]. Maintaining appropriate keratocyte phenotype is vital to preventing them becoming fibrotic, resulting in permanent scar tissue formation [16] [17]. The presence of keratocytes also benefits in nerve re-innervation via neuro-regulatory factor secretion [18]. For these reasons, a limited number of studies have been undertaken to assess the effect of recellularizing the decellularized stroma prior to implantation [10]. In one rabbit host study [10], the porcine scaffold that had been recellularized by injection of human stromal cells and 3 days of culture recovered transparency faster and achieved a more native-like ultrastructure than a decellularized counterpart. In a 12 month long-term study in dogs, decellularized porcine corneas that had been recellularized with human epithelial and stromal cells for 8 days showed improved re-innervation, epithelial integrity and a more normal central corneal thickness, compared with cell-free scaffolds [19].

In acellular clinical studies, where decellularized porcine scaffolds were implanted into the human eye to repair damage from a localised infection, there was a beneficial outcome, without rejection, but with only gradual recovery of transparency and some improvement of visual acuity [1] [4] [3]. The repopulation of the scaffold with keratocytes was slow, as was the healing response compared with using a healthy donated human cornea [1]. In further studies aimed at overcoming such disadvantages, stroma repopulation has been investigated; thin sections of human decellularized scaffold were implanted into patients with advanced keratoconus, but no clear advantage was found between the use of acellular implants compared with those pre-seeded for the short-term of 1 day with autologous adipose mesenchymal stromal cells [5].

Overall, the clinical studies confirm the value of using porcine corneal scaffold, but the potential benefit of re-seeding keratocytes prior to implantation is not clearly established. The first aim of the current study was therefore to examine the *in vitro* recellularization of a decellularized porcine corneal scaffold with human keratocytes over an extended culture period of 28 days and analyse the normality of the resultant cell phenotype. The premise was that, normality established prior to implantation would best result in post-implant transparent tissue. Human keratocytes were chosen for this investigation as these cells have already shown utility with this model of decellularized porcine scaffolds in the rabbit and offer the most potential for clinical translation [10]. The immunological and healing response in the living host can have a profound effect and there is a need to understand if a native phenotype *in vitro* leads to a transparent cornea after implantation. The second aim was therefore to test if a high cell density recellularized scaffold is superior to a decellularized scaffold using an anterior lamellar keratoplasty (ALK) model in the rabbit.

## 2 Materials and Methods

### 2.1 Isolation and decellularization of porcine corneal lenticules

Porcine eyes were collected a few hours after animal death from an abattoir (Rosderra Meats, Edenderry). The eyes were transported while refrigerated and processed within 12 hours. To process the eyes, they were first trimmed and then decontaminated in 2% iodine solution (Videne, Ecolab, Belgium) in sterile phosphate buffer saline (PBS) for 1 minute with gentle agitation and then washed twice in PBS. The lenticule produced had a smooth, dull, convex, apical surface.

Decellularization was performed as previously described [20] using chemicals purchased from Sigma-Aldrich (Arklow). Corneal lenticules were immersed in decellularization solution containing 0.5% (w/v) sodium dodecyl sulphate (SDS) and 1% (v/v) Triton X-100 in deionised water. To promote removal of cells and denatured proteins, samples were placed in an orbital shaker for 72 hours at ambient temperature, changing the solution every 24 hours. The lenticules were then treated with 10 U/mL of both RNAse and DNAse (Sigma) in 10 mM MgCl_2_ solution for 1 hour at 37 °C and tube-rotated at 15 RPM. Next, they were washed and rotated with 200 U/mL penicillin and 200 µg/mL streptomycin (Gibco, ThermoFisher, Dublin) in PBS for another 72 hours, changing the solution daily. They were stored at 4 °C in PBS containing the antibiotics until required and, prior to use, equilibrated in culture medium for 24 hours in a humidified incubator at 37 °C with 5.0% CO_2_; the method of incubation used throughout the studies.

### 2.2 Scaffold transparency and light transmittance

Light transmittance across the visible spectrum from 350 to 700 nm wavelength was quantified using a spectrophotometer (BioTek Synergy HTX, ThermoFisher) with the scaffold immersed in ultrapure water. Glycerol has been shown to re-establish the thickness and curvature of a swollen cornea [21]. For this reason, the scaffold transmittance was then assessed after immersion in glycerol for 2 hours. Following glycerol treatment, optical quality was also evaluated, by holding the scaffold in a transparent petri dish and then viewing printed text through the base [22].

### 2.3 Scaffold recellularization

Human corneo-scleral rims remaining after keratoplasty were supplied from the Irish Blood Transfusion Service Eye Bank, Dublin after appropriate consent of the donor or next-of-kin. Stromal keratocytes were isolated by explant culture [23]. To ensure the cells were consistent across the experiments, they were expanded from cryopreserved stocks and used at passage 4 or 5. Expansion medium consisted of low-glucose Dulbecco’s Modified Eagle’s Medium (DMEM) (HyClone, Fisher Scientific, Dublin) supplemented with 10% foetal bovine serum (FBS), penicillin (100 U/mL) and streptomycin (100 mg/mL) (Gibco).

Several methods are available to recellularize a scaffold [11]. Simple seeding on the surface was chosen as the recellularization method in the present study. Recellularization was carried out in a 12-well tissue culture plate (Cellstar, Greiner, Austria). A polytetrafluoroethylene (PTFE) disk was used to cover the tissue culture surface to prevent cells attaching to the plate. The scaffolds were recellularized by seeding 0.1 x10^6^ stromal cells in 10-15 µl of expansion medium onto the upper surface of the scaffold and allowing cell attachment for 30 minutes of incubation. Scaffolds were then inverted, and a similar application made to the reverse side. Finally, 0.3×10^6^ cells were added in 1 mL of expansion medium and cultured with incubation for 14 days, with a change of media every second day.

In the normal cornea, keratocytes are in general quiescent, that upon injury to the cornea are stimulated into repair phenotypes, but a fibrotic response can lead to scarring and loss of corneal clarity [24]. To investigate the phenotype of stromal cells introduced into the scaffold, three different culture conditions were examined. In the first group (the short-expansion group), culture was for 14 days in the serum-supplemented media already described. In the second group, culture was for 28 days in serum-supplemented medium (the long-expansion group). For the third group, the cell seeded scaffolds were cultured for 14 days in the serum-supplemented media and then switched for a further 14 days, (the keratocytic-expansion group), to a serum-free keratocyte medium composed of DMEM/F12 medium (HyClone, ThermoFisher) supplemented with 50 µg/mL ascorbic acid (Sigma) and 1x insulin-transferrin-sodium selenite (Gibco), giving a final concentration of insulin 10 µg/mL, transferrin 5.5 µg/mL and sodium selenite 6.7 ng/mL. This medium has been shown to stabilise the cells into a more native corneal phenotype [25]involving self-assembly of cell distribution on the scaffold [23]. Scaffolds that were to be implanted decellularized, were treated for the same time and keratocytic-expansion media, without cells.

Limbal epithelial cells were isolated by cutting the remaining donor limbal tissue into quarters and incubating with 2.5 mg/mL dispase (Life Technologies, ThermoFisher) for 1 hour. The cells were removed by scraping the limbal surface with a scalpel blade, then pooled, and centrifuged at 170 G for 5 min. They were then resuspended in epithelial medium and 15 µl placed directly onto the scaffold to give a density of 10^5^ cells/cm^2^. The epithelium medium was composed of a 3:1 mixture of DMEM (Hyclone, ThermoFisher) combined with Ham’s F12 medium (Gibco) and supplemented with 10% FBS (Gibco), 5 µg/mL human recombinant insulin, 0.4 µg/mL hydrocortisone, 2 pM triiodo-L-thyronine, 10^−5^ M isoprenaline hydrochloride, 5 µg/mL transferrin, 180 µM adenine (all Sigma), 10 ng/mL human epithelial growth factor (Source BioScience, Nottingham, UK) and 2 mM L-glutamine, 100 U/mL penicillin and 100 µg/mL streptomycin (all Gibco).

### 2.4 Histology, immunostaining and imaging

Scaffolds were fixed in 4% paraformaldehyde in PBS for 30 minutes, permeabilized with 2% FBS and 0.5% Triton X-100, then stained with 4’,6-diamidino-2-phenylindole (DAPI) and phalloidin-conjugated to tetramethylrhodamine, to visualize cell nuclei and the actin cytoskeleton, respectively. Afterwards, the scaffolds were embedded in paraffin wax and 6 µm thick slices were cut using a microtome (RM2125, Leica Microsystems, UK) and mounted on glass slides (Superfrost plus, ThermoFisher). Native porcine corneas and decellularized scaffolds were stained by three methods: Haematoxylin and eosin (H&E), alcian blue and picrosirius red, to distinguish cells, glycosaminoglycans (GAG) and collagen, respectively.

Where antibody staining followed paraffin wax embedding, antigens were retrieved by heat-mediated citrate buffer. Primary antibodies were: Aldehyde dehydrogenase ALDH1A1 (Ab9883) and ALDH3A1 (Ab76976) from Abcam (Cambridge, UK). Both were used as the ALDH3A1 is a rabbit sourced antibody, making it unsuitable as a primary in the rabbit, and also, unlike humans, rabbits primarily express ALDH1A1 in the cornea, with negligible amounts of ALDH3A1 [26]; alpha smooth muscle actin, α-SMA (Ab7817); fibronectin (Ab6328) from Abcam; collagen Ia (sc-59772) from Santa Cruz Biotechnology (Heidelberg, Germany); collagen III (GTX26310) from GeneTex (Insight Biotechnology, London, UK); and keratocan (HPA039321) from Atlas Antibodies (Cambridge Bioscience, Bar Hill, UK). Appropriate secondary antibody conjugated with AlexaFluor 488 (Abcam) was visualised on a laser scanning confocal microscope (SP8, Leica). Negative controls involved use of the secondary antibody alone. Cross-reactivity of a nucleolar stain for human cells meant that we were unable to confirm the survival of human cells after the rabbit ALK.

The density of the scaffold limited imaging to less than 100 µm from the surface. To assess the extent of cell position in the deeper layers, for each sample, 3 scaffold cross-sections were cut, and every DAPI-labelled nucleus was classified as a cell in the section. The closest scaffold surface was assumed to be the origin of each cell, and the distance from that surface to the nucleus was measured for each. Cells were grouped into 20 µm depths and counted to quantify the extent of cell penetration.

### 2.5 Biochemical analyses

Samples were held frozen at −80 °C prior to biochemical analysis. All weights were standardised against the dry weight of the sample. For the DNA, glycosaminoglycans and collagen quantification, native lenticules and decellularized scaffolds were freeze-dried then weighed. The freeze-dried samples were then digested in 3.88 U/mL of papain solution and tube-rotated at 60°C for 18 hours to disrupt the tissue.

#### 2.5.1 DNA content

The DNA was fluorescently stained using a Quant-iT PicoGreen double strand (ds) DNA kit (Molecular Probes, Biosciences, Dublin) and quantified against a control standard by spectrophotometry at 480 nm excitation and 520 nm emission wavelengths. The extent of DNA fragmentation was also measured after extraction of the DNA using a DNeasy kit (Qiagen, ThermoFisher) and separation of the fragments by electrophoresis on a 3% agarose gel.

#### 2.5.2 Glycosaminoglycans content

Determination of GAG used a 1, 9 dimethylmethylene blue dye assay (Blyscan, Biocolor, UK). Briefly, 10 µl of papain digested sample was incubated with the dye for 30 minutes at ambient temperature with tube-rotation, then centrifuged at 15,000 G for 10 minutes and the supernatant removed. The remaining pellet was treated with the dye dissociation reagent and the dye density measured with a spectrophotometer at 656 nm excitation wavelength. Known quantities of GAG were processed in the same way to obtain a dye absorption standard curve.

#### 2.5.3 Collagen content

The collagen content was determined using a hydroxyproline assay [27] which is based upon the fixed ratio of hydroxyproline to collagen of 1 to 7.69 [28]. To measure the hydroxyproline, a 10 µl of papain digested sample was dissociated with 38% HCl at 110 °C for 18 hours and then dried for 48 hours at 50 °C. Serial dilutions of trans-4-hydroxy-Lproline (Fluka, ThermoFisher) in papain solution were used to obtain a standard range. Samples and standards were incubated for 20 minutes at ambient temperature with n-propanol buffer and chloramine-T to oxidize the hydroxyproline and stained with a 2,4-dimethoxybenzaldehyde dye for 20 minutes at 60 °C. The dye concentration was quantified at 570 nm excitation wavelength using a spectrophotometer.

#### 2.5.4 SDS contamination quantification

Following decellularization, the extent of SDS contamination of the scaffold was measured by a methylene blue assay, whereby the complex of the SDS anion and dye cation is quantified by dye light-absorption. Freeze-dried scaffolds were weighed, vortexed in 1 mL of deionised water and then macerated for six hours. A 250 μl sample was then taken and vortexed with an equal volume of 250 μg/mL methylene blue solution (Sigma-Aldrich), and 1 mL of chloroform added. The mixture was centrifuged on a benchtop centrifuge to separate the aqueous supernatant. The absorbance of the chloroform and dye infranatant was measured at 665nm using a plate reader (BioTek Synergy HTX). Absorbance was quantified against a standard of deionised water and serial dilutions of 0.5% w/v SDS.

### 2.6 Gene Expression

Trizol (Invitrogen) was used to extract RNA from cells in the scaffold in combination with homogenisation (IKA T10 basic, Sigma) [29]. Chloroform was then added and vortexed, prior to centrifugation at 4 °C for 15 minutes at 12,000 G to remove debris. To precipitate the RNA, the supernatant was transferred into a new tube and diluted with the same volume of isopropanol and 3 µl of Glycoblue (Life Technologies, ThermoFisher). After overnight freezing to −20 °C, centrifugation was repeated, and the supernatant discarded. The pellet was washed in 70% ethanol in RNAse-free water, centrifuged, the supernatant removed, air-dried, then dissolved in RNAse-free water.

RNA yield and purity were measured using a NanoDrop-1000 (ThermoFisher). A high capacity cDNA reverse transcription kit (Invitrogen) was then used to convert RNA into cDNA. Real-time PCR was performed using TaqMan Universal Master Mix II (ThermoFisher) along with the following TaqMan primers: glyceraldehyde-3-phosphate dehydrogenase (GAPDH, Hs02758991_g1), aldehyde dehydrogenase 3A1 (ALDH3A1, Hs00964880_m1), alpha smooth muscle actin (α-SMA) (ACTA2, Hs00426835_g1), keratocan (KERA, Hs00559942_m1), collagen I (COL1A1, Hs00164004_m1), lumican (LUM, Hs00929860_m1), decorin (DCN, Hs00754870_s1) and CD34 (Hs00990732_m1). The genes of interest were normalized against glyceraldehyde 3-phosphate dehydrogenase using the ΔΔC_t_ method. Calculated values were expressed as a power of 2^−ΔΔCt.^ All values were normalized against the serum-expanded cells growing in tissue culture prior to seeding onto the scaffolds.

### 2.7 Axonal outgrowth test

Dorsal root ganglion (DRG) have been used to validate the ability of nerves to innervate a collagen hydrogel scaffold *in vitro* which was subsequently confirmed in a 12 month animal model [30]. To perform a similar *in vitro* test, DRGs were isolated from adult rats, rinsed in expansion media and placed directly onto scaffold that had been removed from its expansion medium, but left moistened. The ganglia were left for 1 hour of incubation and then 1 mL of DMEM, supplemented with 10% FBS and 10 ng/mL NGF (Sigma), was added to cover it. Media was changed every 3 days. After 14 days in culture, scaffolds were fixed in paraformaldehyde and immunostained with the neural cell marker βIII-tubulin (Sigma).

### 2.8 *In vivo* scaffold implantation in rabbits

Eight female New Zealand rabbits (21-23 weeks old, 2.75-4.0 kg) were used for the experiment, randomly divided into 2 groups. Additional rabbits were used to establish the model. The selection of decellularized or recellularized samples was randomised and the surgical and histological examination teams were unaware of the implant cellularization status until the data collection was complete. This resulted in 3 acellular and 5 cellular implants. Animal welfare and surgery procedures were approved by Research Ethics Committees of both Trinity College Dublin (301107-1001) and Dublin City University (DCUREC/2018/253) and the project authorised by the Health Products Regulatory Authority (AE19136/PO93). The right eye of each rabbit was used for the anterior lamellar keratoplasty, whilst the left eye was untreated, as a control. One animal group received decellularized porcine scaffold left acellular, and the other group, scaffold that had been recellularized with human stromal cells. Both had received the keratocytic-expansion.

As detailed elsewhere [31], prior to surgery the rabbits were first pre-medicated with subcutaneous buprenorphine (0.03 mg/kg) (Buprecare, Animalcare Group, York, UK) and then anesthetized with intravenous ketamine (10 mg/kg) (Hameln pharmaceuticals Gloucester, UK) and xylazine (3 mg/kg) (Sedaxylan, Dechra, Shrewsbury, UK) whilst oxygen was administered through a face mask to maintain oxygen saturation. To extend anaesthesia, 0-5% gaseous isoflurane (Isoflurin, Vetpharma Animal Health, Barcelona, Spain) was used and additional intravenous ketamine and xylazine given as a bolus, as necessary.

After the epithelium had been debrided with 70% ethanol over the implant site, the recipient cornea was trephined to 250 µm depth using a 6.0 mm diameter Barron radial vacuum trephine (Barron Precision Instruments, Grand Blanc, MI, USA) and the stromal tissue excised by the same method as with the porcine tissue. Scaffolds were orientated, convex side apical, to maintain the native density of the scaffold collagen fibrils. Scaffolds were sutured into the rabbit defect with 16 interrupted 10–0 nylon sutures (Serag Wiessner, Naila, Germany or Ethicon, Medray, Dublin) and the knots buried to reduce irritation. Immediately after surgery the eye was irrigated with the broad-spectrum antibiotic, cefuroxime (Aprok, Biopharma S.R.L., Roma, Italy).

Under the control of a veterinarian, postoperative analgesia of subcutaneous buprenorphine (0.01 mg/kg) was administered twice a day for at least 3 days. Postoperative topical corticosteroid, (Betnesol 0.1% w/v, RPH Pharmaceuticals AB) drops or ointment, and topical antibiotic chloramphenicol drops (Minims chloramphenicol 0.5% w/v, Bausch Health, Dublin) or ointment (1% w/w, Martindale Pharma, UK), were applied up to four times daily.

Post-operative eye examinations were performed at: 1, 3, 7, 14 days and 1, 2 and 3 months. At these times, slit lamp biomicroscopy (Portable slit lamp, Reichert, Buffalo, NY, USA) was used for the general assessment of the health of the anterior segment of the eye and intraocular pressure was monitored by contact tonometry measurement (Tonovet plus, Icare, Vantaa, Finland). The extent of re-epithelialization of the implants was confirmed by fluorescein staining (Minims fluorescein sodium 1%, Bausch Health). At 3 months post-operative, the rabbits were euthanized with an overdose of intravenous sodium pentobarbital (Dolethal, Vetoquinol, Towcester, UK). Eye globes were removed, photographed and examined by optical coherence tomography (OCT) with an OC LabScope (Lumedica, Durham, NC, USA). They were then fixed in 10% formalin for 24 hours at ambient temperature, the cornea was then excised and embedded in paraffin wax for histological and immunohistochemical analysis.

### 2.9 Statistical analysis

All experiments were performed three times with a minimum of three replicates. Data are presented as the mean ± SD. Statistical analyses were calculated using GraphPad Prism 6 (GraphPad Software, San Diego, USA). An unpaired two-tailed *t*-test was used to compare two groups and a 2-way ANOVA and Tukey post-hoc analysis when comparing three or more groups. Statistical significance was accepted at a level of p < 0.05.

## 3 Results

### 3.1 Decellularization

Decellularization requires cell removal and DNA minimisation without marked removal of other constituents or the addition of chemical contamination. DAPI and H&E staining demonstrated that the cellular structure of native tissue (Fig 1A) was removed by decellularization and no nuclei could be seen (Fig 1B). Supporting this, Picogreen quantification of DNA showed a reduction from the 728.9 ± 110.2 ng/mg of the native cornea to 14.89 ± 5.56 ng/mg following decellularization (Fig 1C). The display of fraction size of DNA > 1500 base pairs, taken to be supercoiled DNA, was a bright band for the native cornea, while only faint smearing for the decellularized scaffold lanes (Fig 1F). GAGs were also reduced, visualised by the reduction of alcian blue staining (Fig 1A & B) and quantified by the biochemical assay (Fig 1D); in the native cornea GAG was 40.13 ± 2.15 µg/mg, while in the decellularized this was 10.22 ± 5.44 µg/mg.

**Fig 1.**
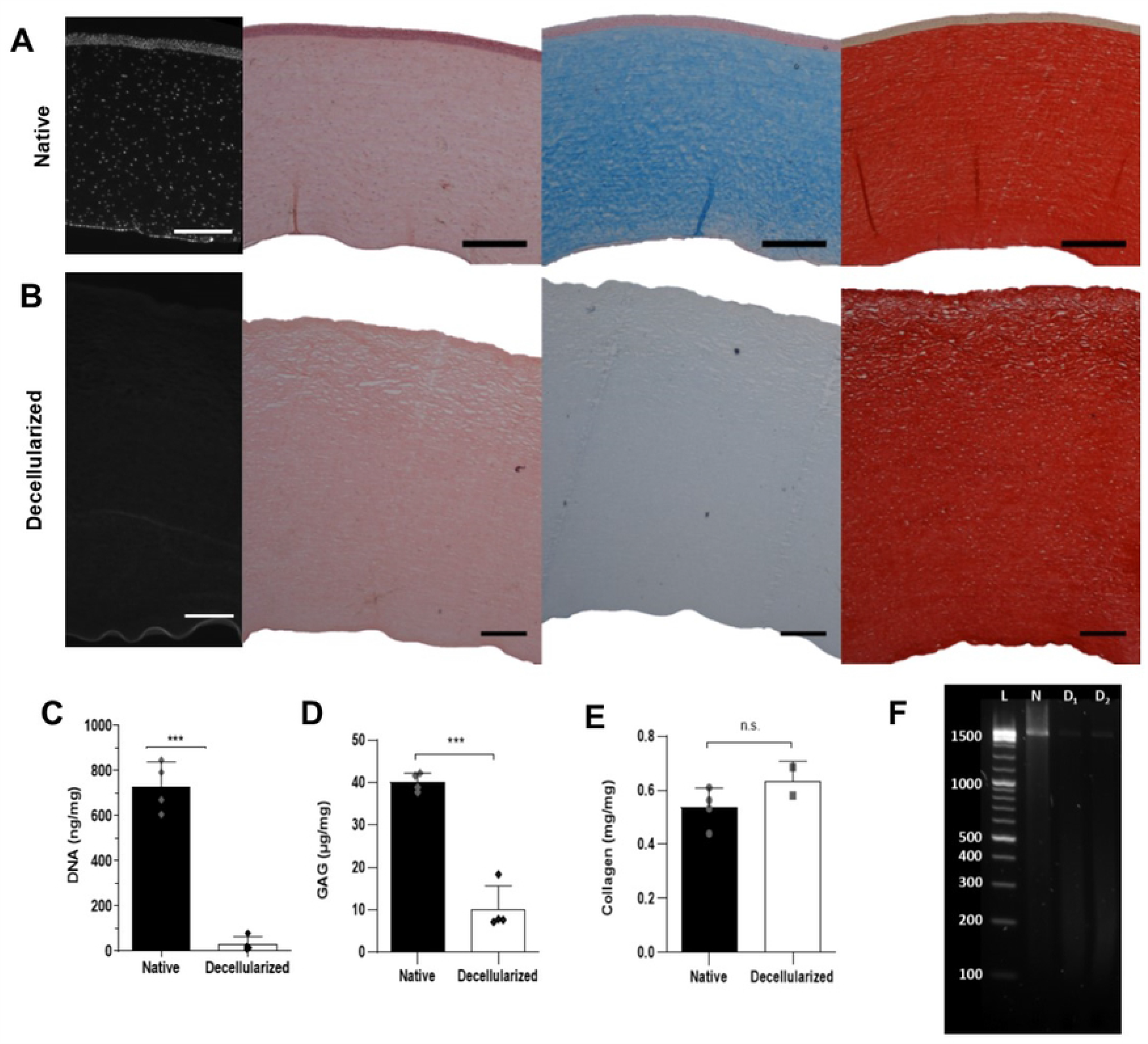
Evaluation of porcine cornea decellularization. A) Native cornea and B) decellularized scaffold, from left to right stained with, DAPI, H&E, alcian blue or picrosirius red. Black scale bar = 500 µm, white = 250 µm. Quantification of native cornea and decellularized scaffold C) DNA, D) glycosaminoglycans and E) Collagen. p * < 0.05, p ** < 0.01, p *** < 0.001. F) Agarose gel electrophoresis of DNA from native cornea (N) and decellularized scaffold (D1 and D2). Ladder (L).

Collagen was not noticeably reduced by decellularization, as seen by picrosirius red staining (Fig 1A & B) and quantified by the hydroxyproline assay (Fig 1E). The collagen content rose from 0.59 ± 0.07 mg/mg to 0.63 ± 0.07 mg/mg following decellularization, as its proportion increased as other constituents were reduced. From the histological images, the collagen structure appeared similar between the groups, other than the collagen in the anterior third of the decellularized scaffold appeared more separated, as if there had been more osmotic swelling resulting in lacunae.

To assess chemical contamination of the scaffold, the concentration of SDS after decellularization was measured to be 52 ± 23 µM.

In PBS the scaffold showed haze, but when dehydrated in glycerol there was good optical clarity (Fig 2). Light transmittance was poor in water (0.95% at 300 and 43.72% at 700 nm), but when transferred to glycerol it improved comparable with that of the native tissue.

**Fig 2.**
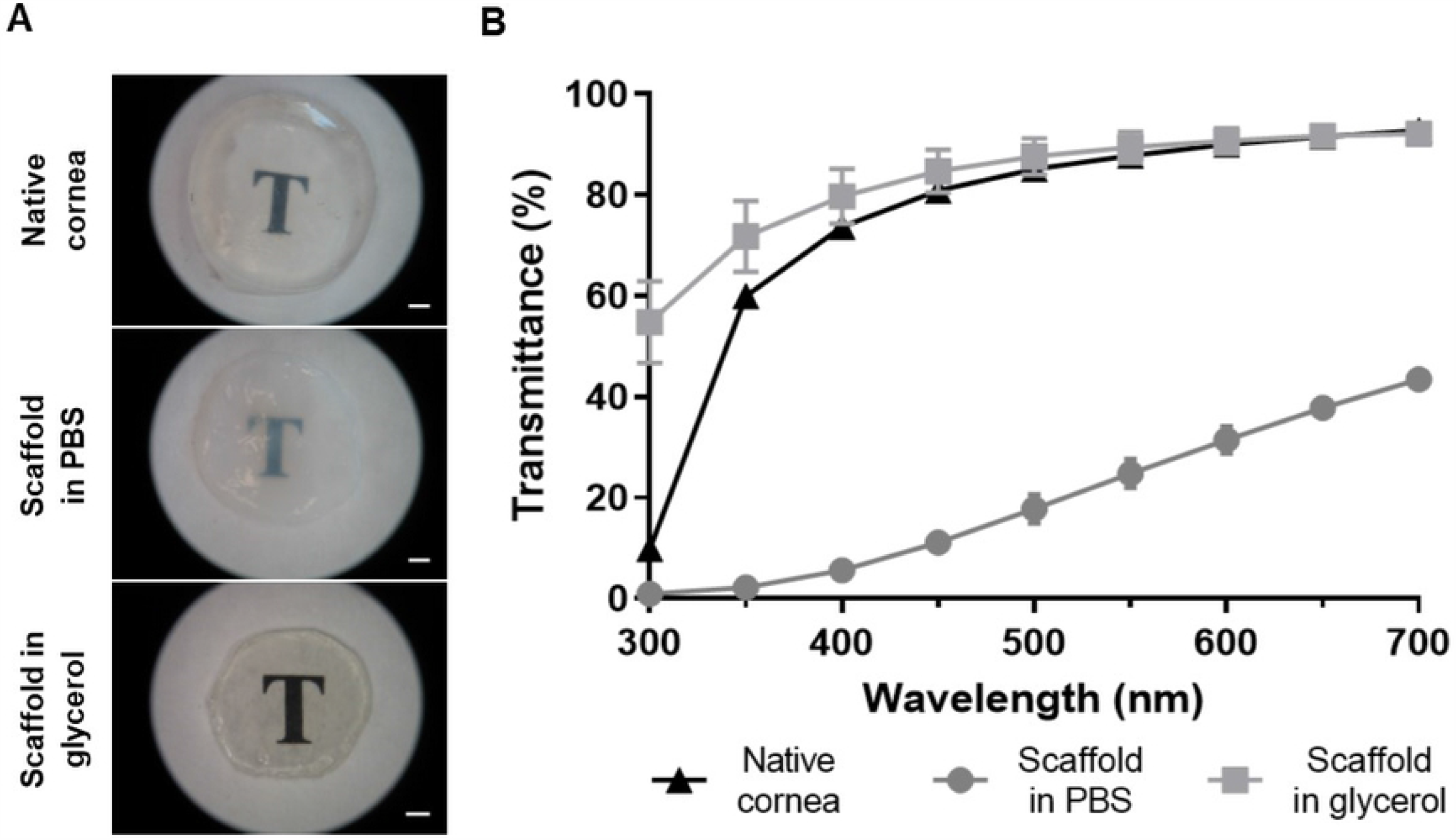
Scaffold optical clarity and light transmission. A) Macroscopic optical clarity. Scale bar = 1 mm. B) Light transmittance spectra.

### 3.2 *In vitro* stromal repopulation

Keratocytes were able to penetrate into the scaffolds regardless of the culture conditions examined (Fig 3A). Confocal microscopy of the recellularized scaffold demonstrated a higher cell density close to the surfaces, diminishing as the depth increased (Fig 3B). The count of cells for every 20 µm of depth in corneal sections confirmed density decreased with depth (Fig 3C) and all three culture conditions had a similar depth/density profile. The median depth was 66 µm for the short-expansion and 71 µm for both the long-expansion and the keratocytic-expansion. Measurement of keratocyte DNA quantity in the scaffold did not vary significantly between the three culture conditions (Fig 3D). Deeper cells had a more native organisation with a dendritic morphology and interconnecting pseudopodia that appeared to form a 3D network.

**Fig 3.**
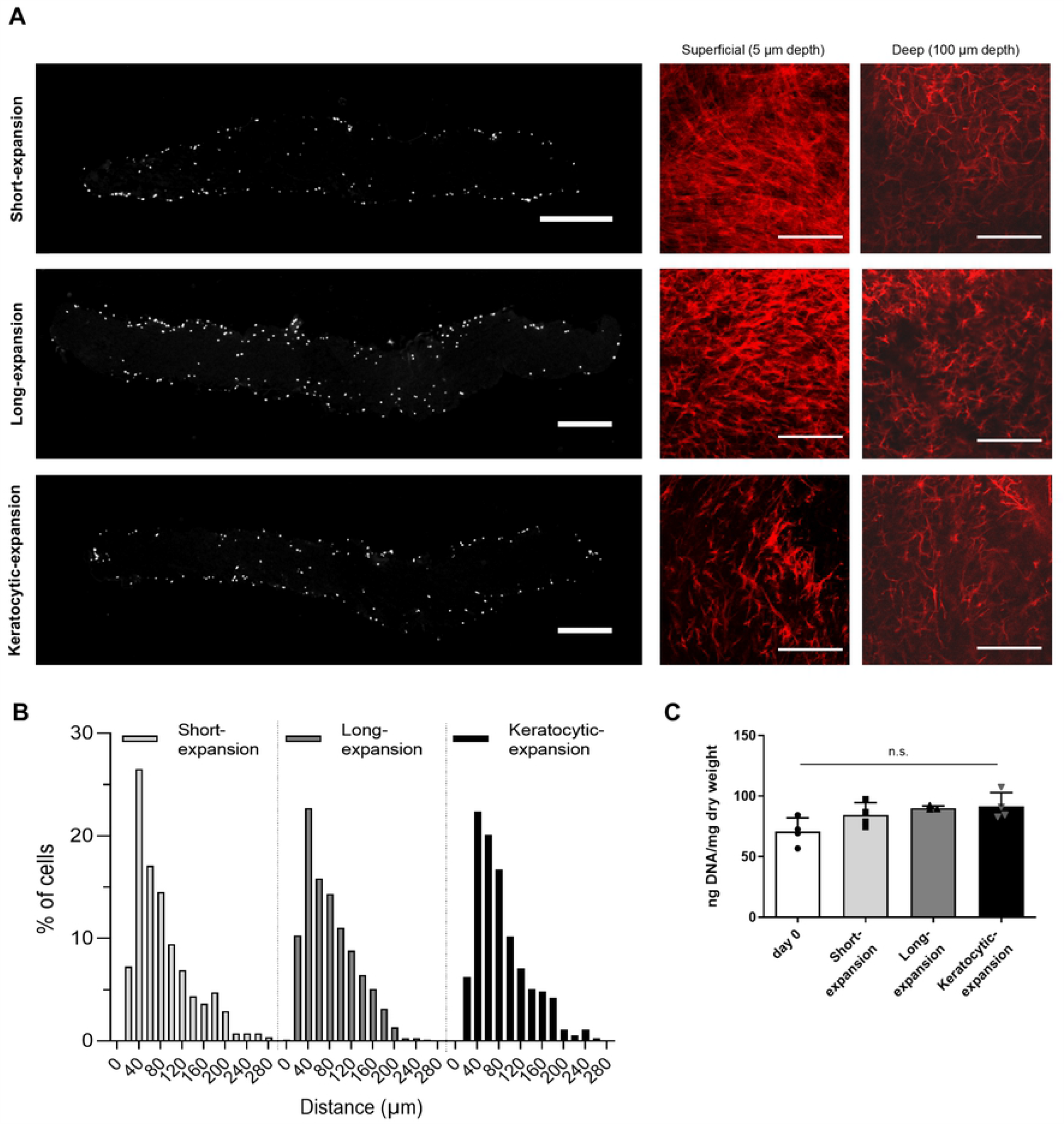
Analysis of the effectiveness of recellularizing scaffold with human keratocytes. A) Longitudinal sections of recellularized scaffolds of keratocytes stained with DAPI nuclear staining. Scale bar = 500 µm. B) Actin staining of keratocytes at a superficial depth (5 µm), and deep depth (100 µm). Scale bar = 200 µm. C) Quantification of number of keratocytes grouped by distance migrated from the surface of the scaffold. D) DNA quantification of recellularized scaffolds (n.s. = not significant).

The gene expression of cells in the repopulated scaffolds was assessed by quantitative PCR (Fig 4). Keratocyte markers for the crystallin ALDH3A1, CD34 and small leucine-rich extracellular matrix proteoglycans keratocan, decorin and lumican, as well as collagen I and the fibrotic marker α-SMA were all significantly up-regulated with the keratocytic-expansion treatment compared with the short- and long-expansions.

**Fig 4.**
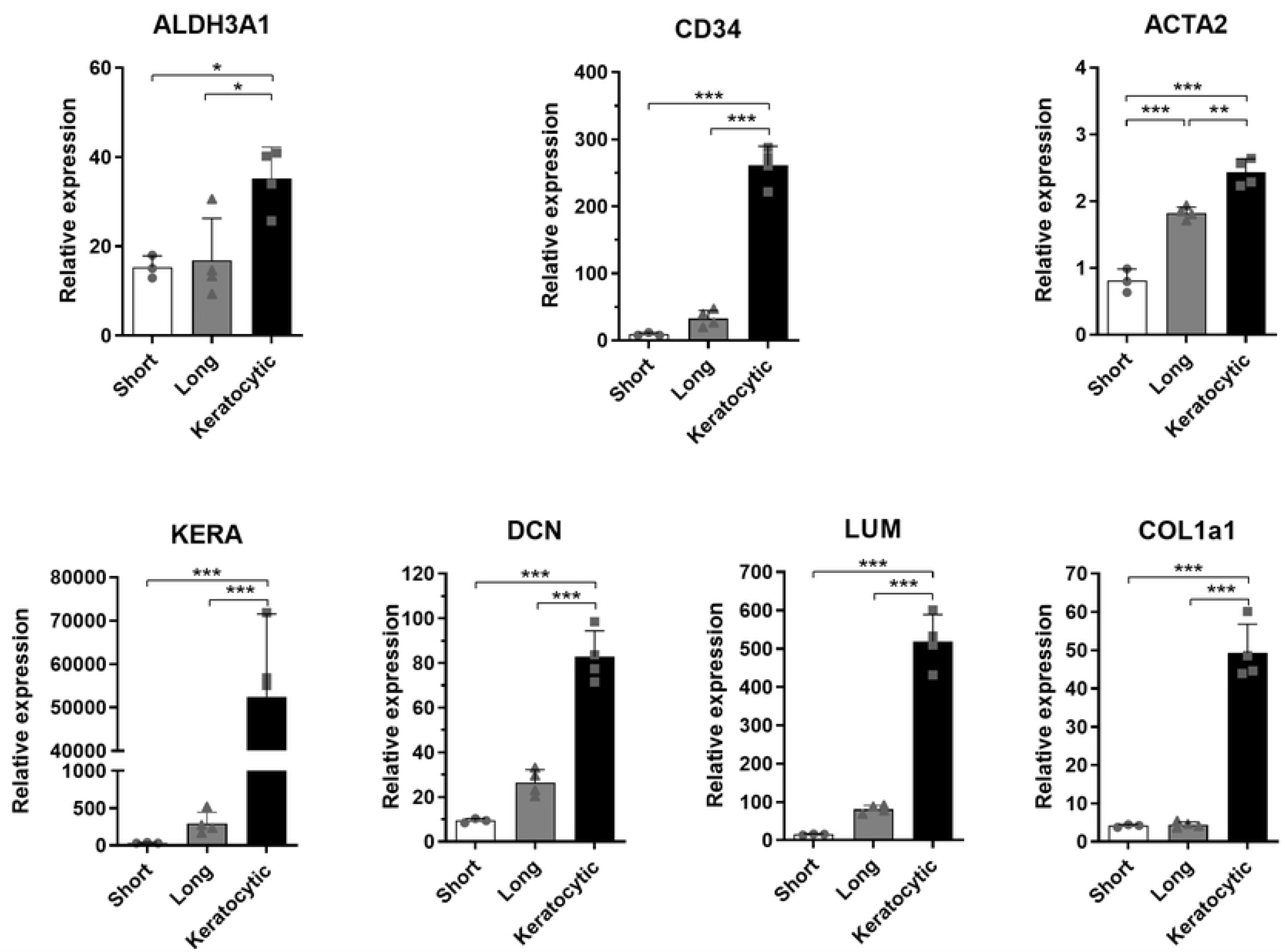
Gene expression analysis of keratocytes in the scaffold. There is a significant difference for all gene expression compared with that of the keratocytic-expansion. (n = 3-4). * p < 0.05, ** p < 0.01, *** p < 0.001.

Cell phenotype was further analysed via immunohistochemistry for ALDH3A1, keratocan and α-SMA (Fig 5) in the scaffold. Native human central cornea and limbus were used as controls. The short-expansion group was negative for all three markers. The long-expansion group and keratocytic-expansion were also negative for keratocan and α-SMA. However, the long-expansion showed faint ALDH3A1 staining and the keratocytic-expansion had much brighter expression indicating greater maturity.

**Fig 5.**
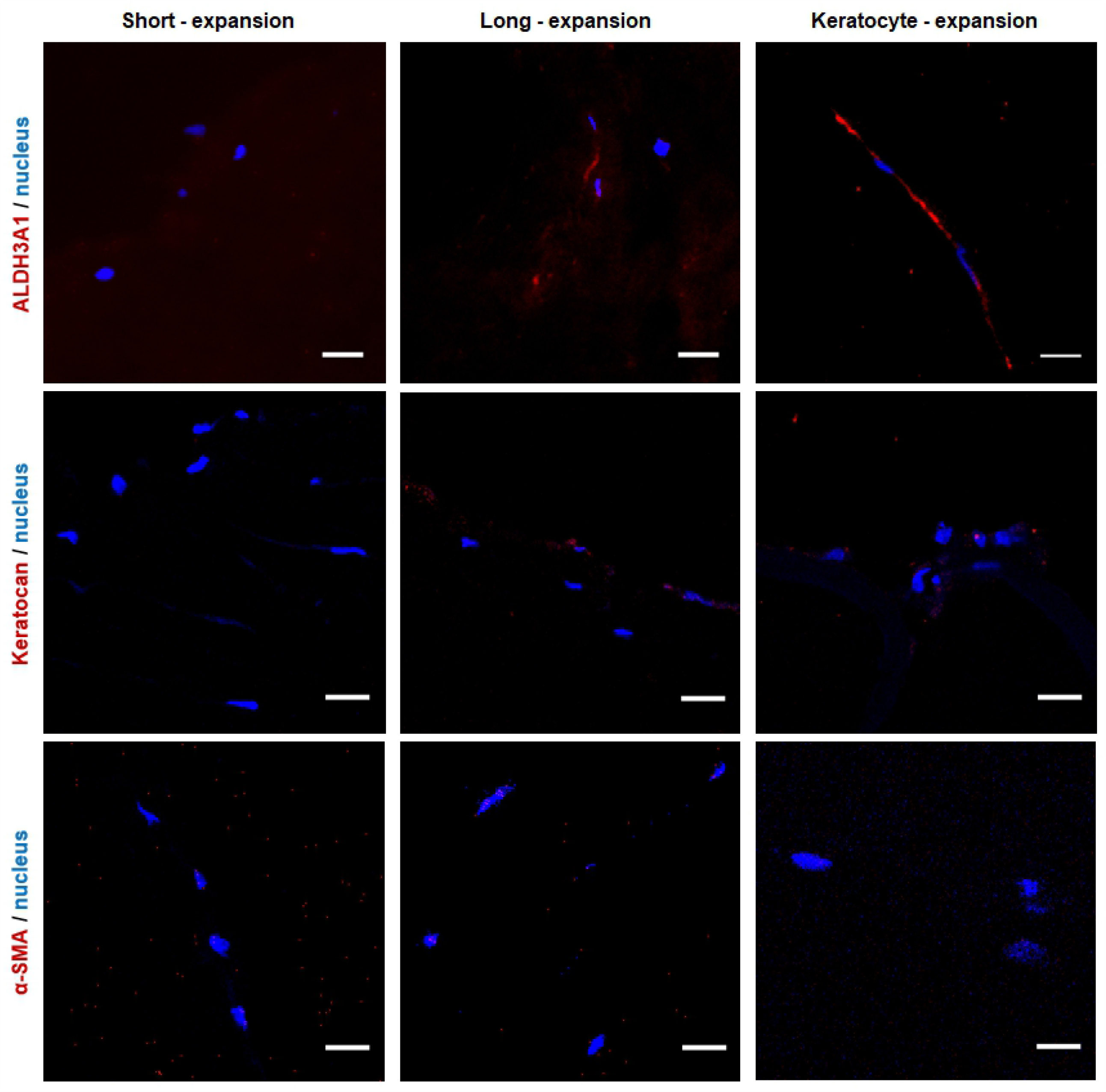
Phenotype of keratocytes in the scaffold. Keratocyte markers, ALDH3A1 and keratocan. Fibrotic marker α-SMA. Nucleus, DAPI nuclear staining (Scale bar = 20 µm).

### 3.3 *In vitro* epithelial growth confirmation

Freshly isolated human limbal epithelial cells colonized the surface when seeded onto the scaffold. There was confirmation of native morphology, as the cells adopted the typical cobblestone appearance with a high nucleus-to-cell size ratio (Fig 6A).

**Fig 6.**
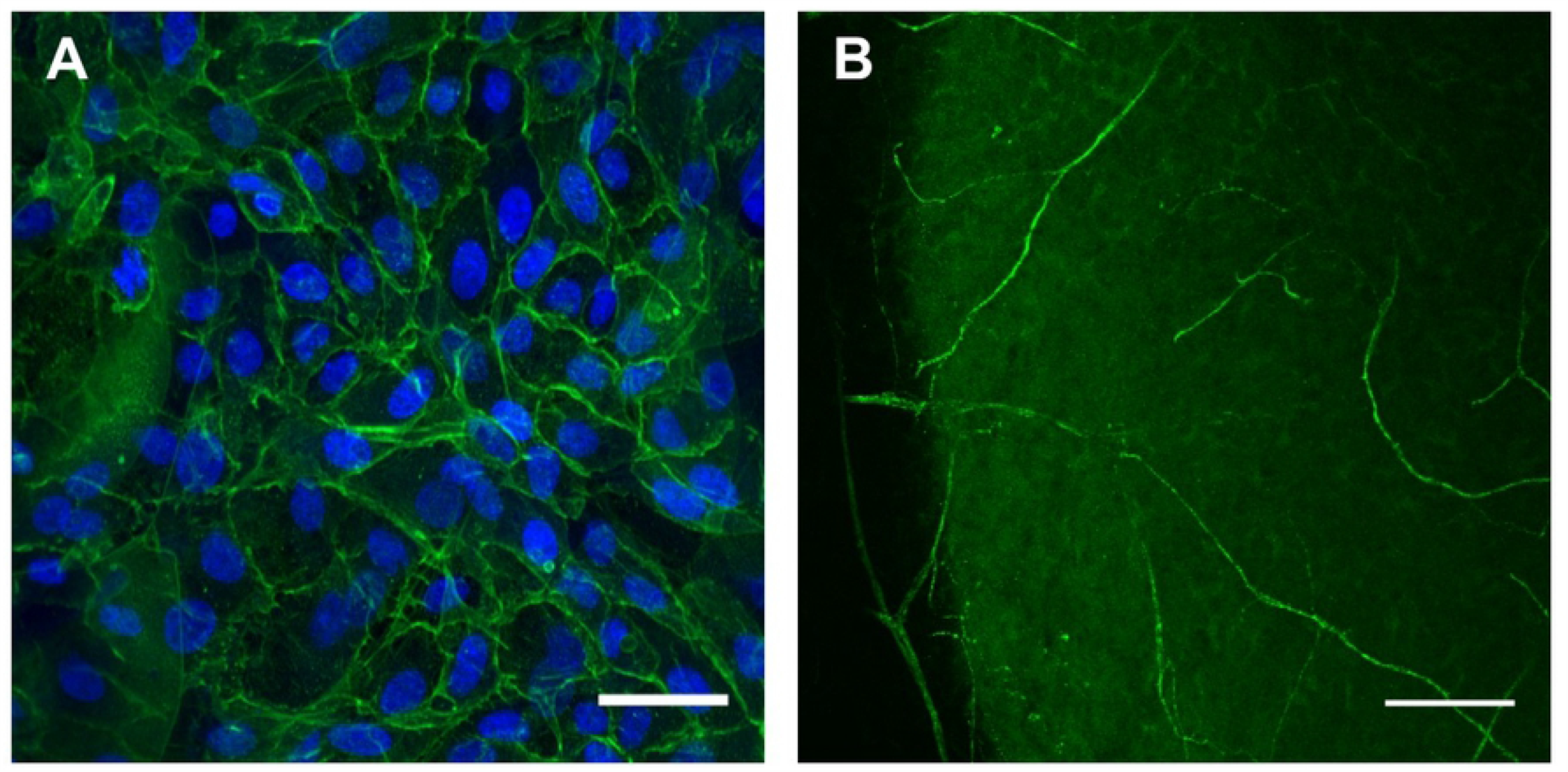
Confirmation of *in vitro* recellularization of porcine scaffold. A) Rabbit corneal epithelial cells. Scale bar = 50 µm. B) Rat DRG βIII-tubulin-positive neurites permeate the scaffold. (Scale bar = 100 µm).

### 3.4 *In vitro* axonal growth confirmation

Axon regrowth into the scaffold for 14 days was demonstrated by βIII-tubulin-positive neurites from the rat DRG extending through to the centre of the scaffold (Fig 6B).

### 3.5 *In vivo* implantation of scaffolds

#### 3.5.1 *In vivo* operative procedure

The scaffolds were successfully implanted using conventional surgical technique (Fig 7A&B). Some scaffolds appeared contracted prior to surgery, but readily relaxed into place. Implants were easily transferred and positioned with conventional surgical instruments whilst displaying excellent suture retention strength characteristics. OCT showed that the defect was repaired with close apposition of the scaffold with the native tissue (Fig 8B).

**Fig 7.**
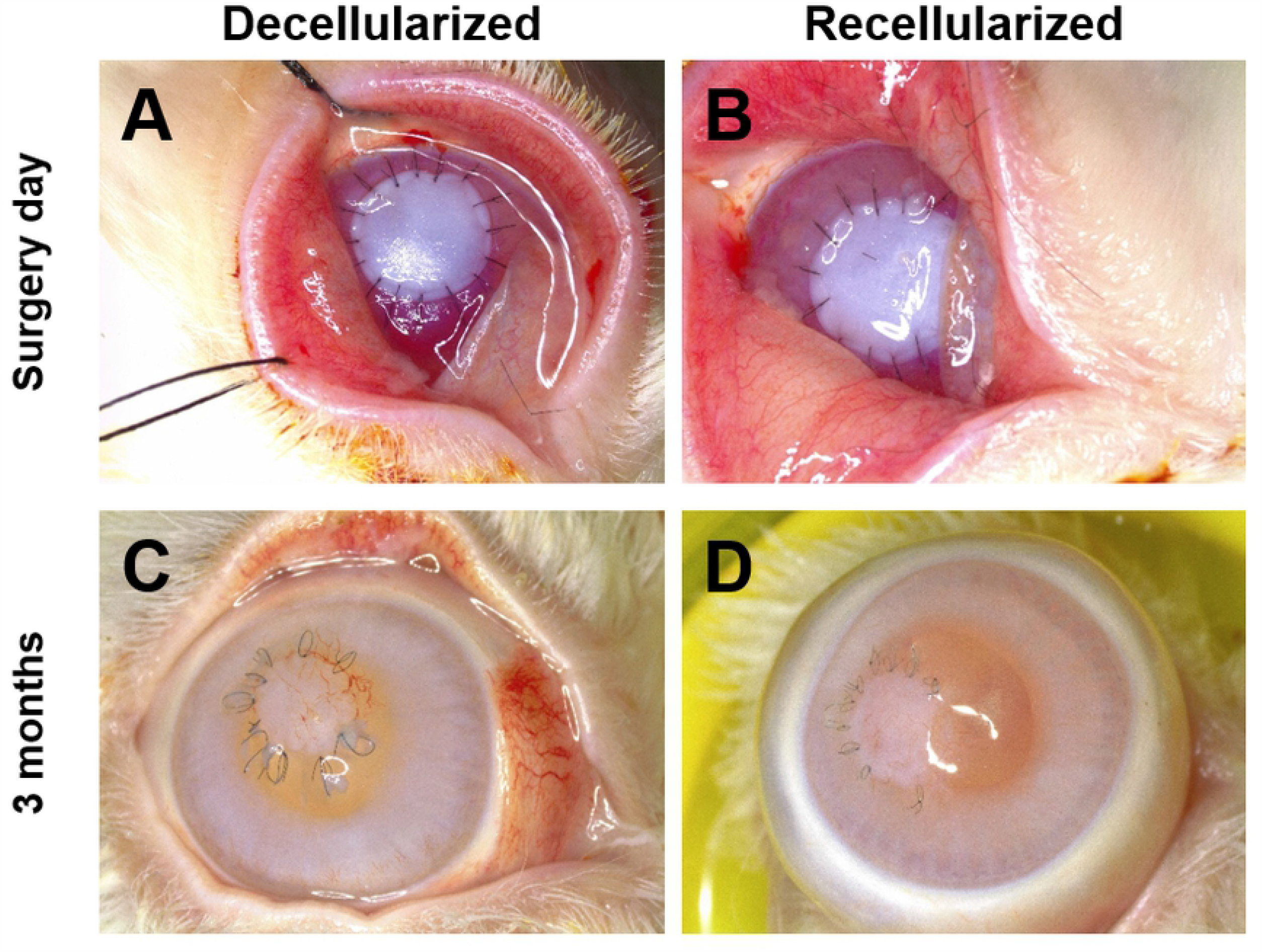
Macroscopic appearance of scaffold in the rabbit eye. Decellularized scaffold (left) with a recellularized scaffold (right). A & B) Day 0. C & D) 3 months post-implantation.

**Fig 8.**
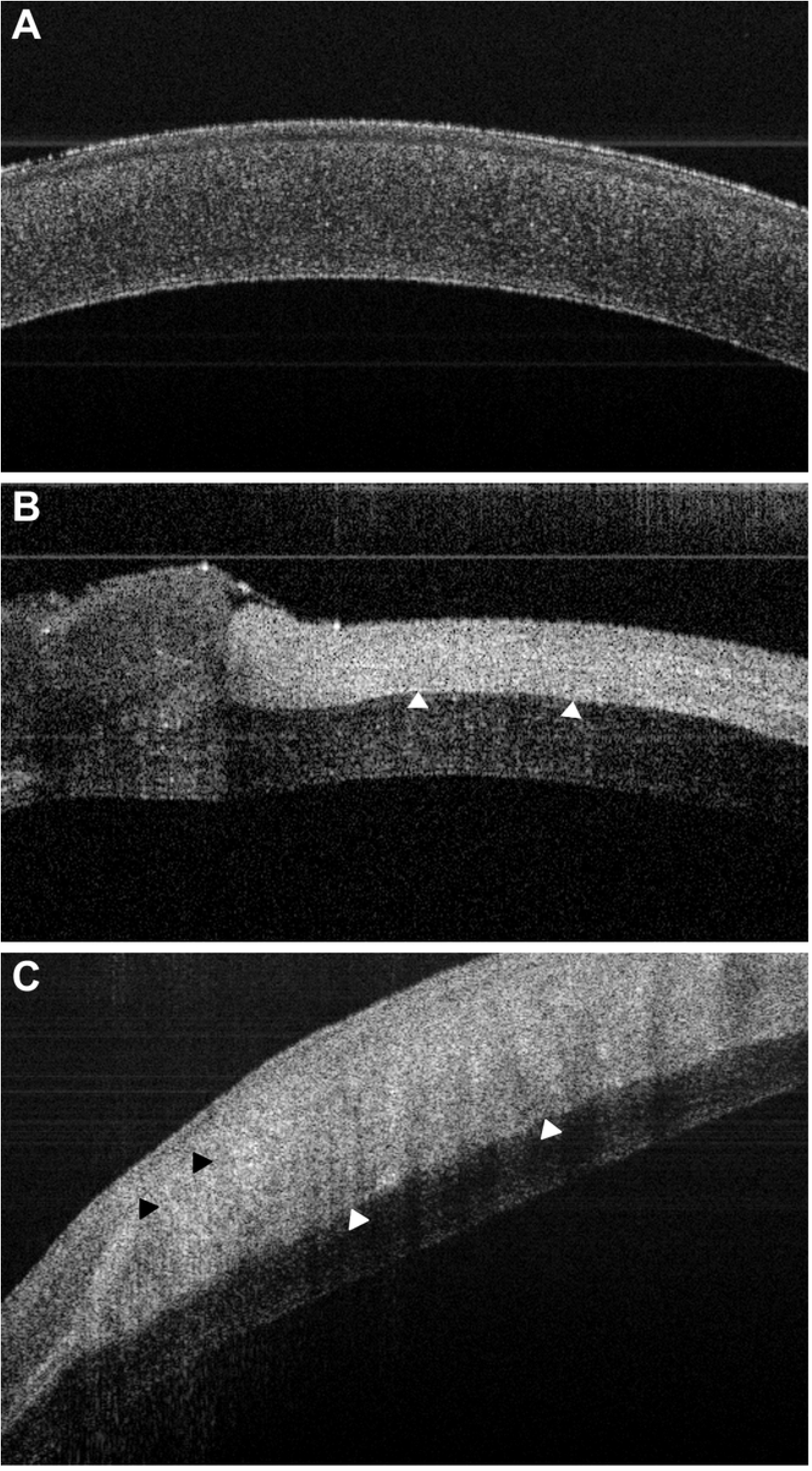
OCT of the rabbit eye. A) Normal un-operated cornea. B) At the time of surgery, day 0. C) 3 months post-surgery. At day 0 the scaffold can be seen to be well-apposed with native tissue (white arrows) repairing a deep defect. At 3 months post-surgery an apparent regeneration area can also be seen anterior to the scaffold (black arrows).

#### 3.5.2 *In vivo* post-operative progress

Fluorescein staining demonstrated regeneration of the epithelium over most of the scaffold surface 3 weeks post-implantation (Fig 9A, B). By 2 months there was complete cover (Fig 9C, D). Over the experimentation period, the scaffolds appeared to decrease in diameter. The sutures became looser, to the point that a few were unburied and were removed for animal welfare. Although the original whiteness of the scaffold did subside, with some areas appearing to be more translucent. Neovascularization developed, but intraocular pressure did not elevate.

**Fig 9.**
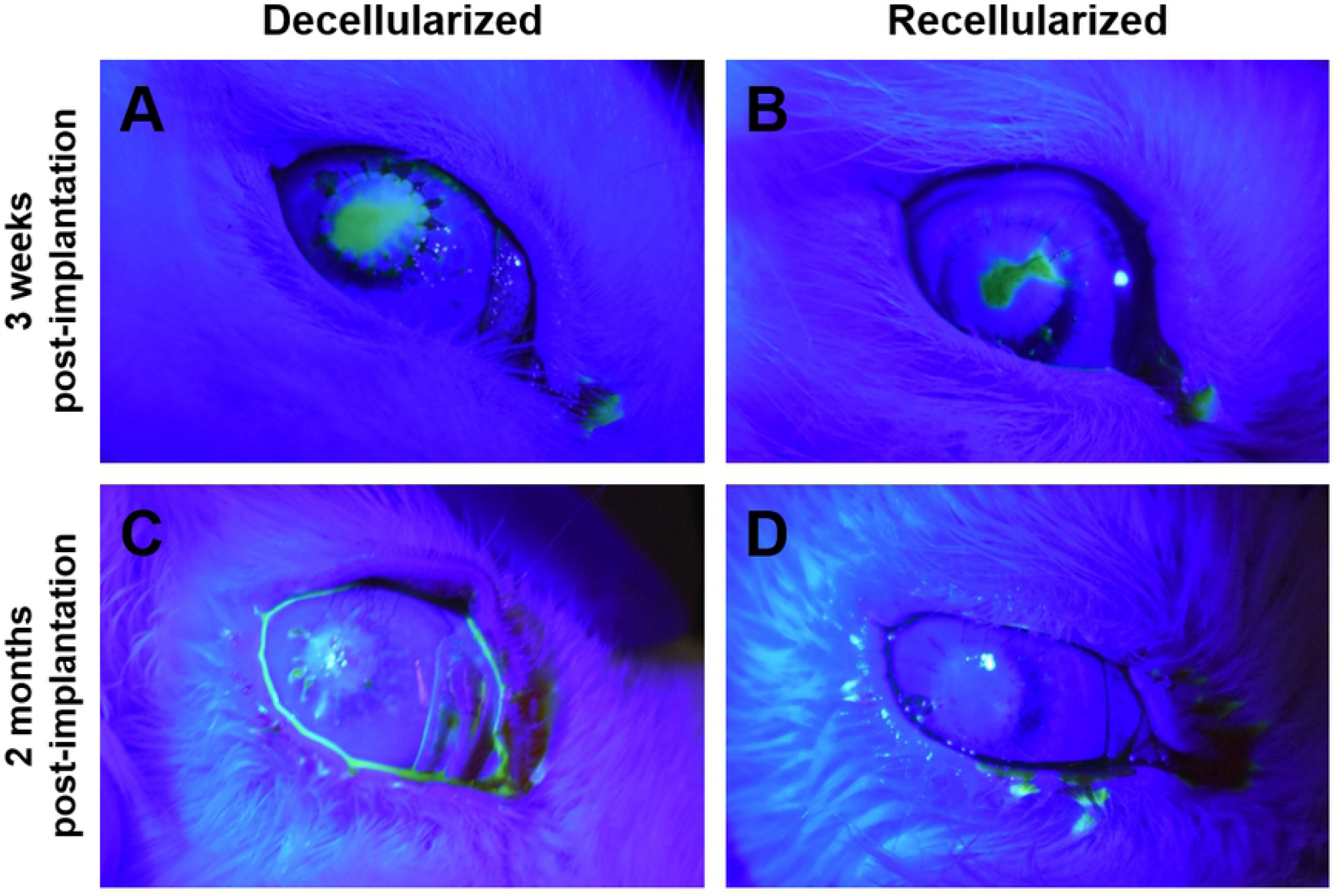
Fluorescein staining of the rabbit eye. A,C) 3 weeks and B,D) two months post-implantation. With both groups the epithelial barrier is not complete at the earlier time, but is intact later.

Three months after implantation, decellularized and recellularized showed comparable epithelial layer re-establishment transparency had not returned to the implanted scaffolds (Fig 7C,D). The location of the opacification appeared to be in the anterior layer.

### 3.6 Examination of post-mortem eyes

The animals were euthanized, and OCT showed that the scaffold remained in place, but it appeared swollen (Fig 8C). Three regions were discerned: a posterior native tissue, a central scaffold and an anterior area. The eyes were removed, and the corneas used for histological analysis, with the unoperated eye as control.

#### 3.6.1 *Post-mortem* corneal ECM appearance

The stroma of both groups gave the appearance of three layers: a posterior native area, a central scaffold area and an anterior regenerating area (Fig 10). The central scaffold area appeared denser than the other two areas when viewed with picrosirius red staining. The scaffold area was strongly positive for collagen I, indicating retention of the implanted collagen in both groups. Collagen III was localised to the apical area and adjacent to the scaffold in the posterior native area; it was not present in the scaffold itself. Similarly, fibronectin was also located in this apical area, particularly with the recellularized group (Fig 10).

**Fig 10.**
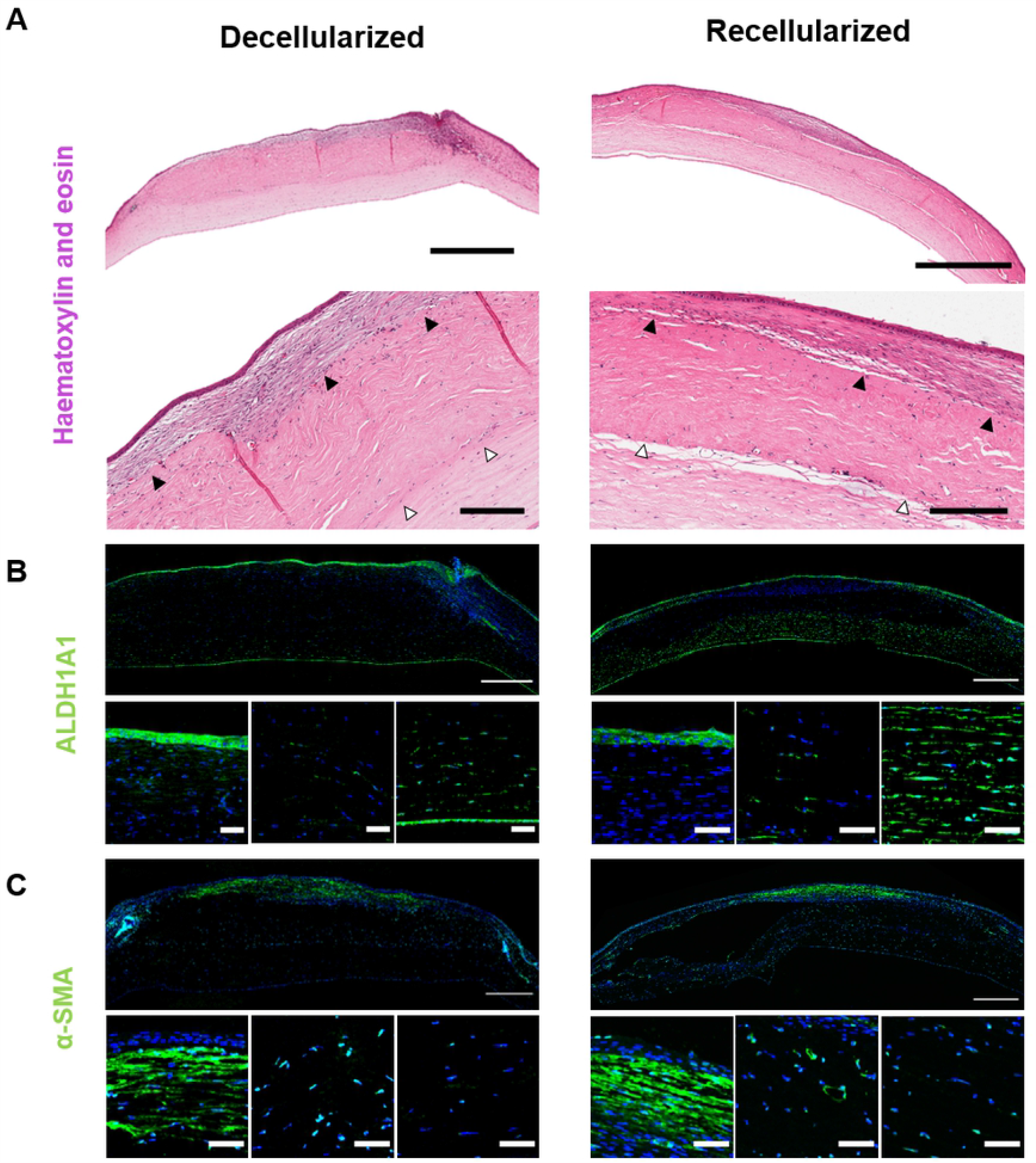
Rabbit cornea 3 months post-implantation stained for ECM content. Representative images comparing a decellularized scaffold (left) with a recellularized scaffold (right). Picrosirius red staining (Scale bar = 200 µm) and immunostaining against collagen I, fibronectin and collagen III, with DAPI nuclear stain (Scale bar = 500 µm). Scaffold apposed with native tissue (white arrows) and regeneration area anterior to the scaffold (black arrows). The Descemet’s layer separation is a processing artefact.

#### 3.6.2 *Post-mortem* corneal cell appearance

H&E staining showed that there was a complete epithelial covering of all samples. This was at least 2 to 3 cells deep (Fig 11A). There was a sparse, apparently normal, distribution of cells in the central scaffold and posterior native area of both groups (Fig 11A,B). The anterior area appeared to be regenerating, with an increased density of cells. The corneal endothelium was intact in all cases.

**Fig 11.**
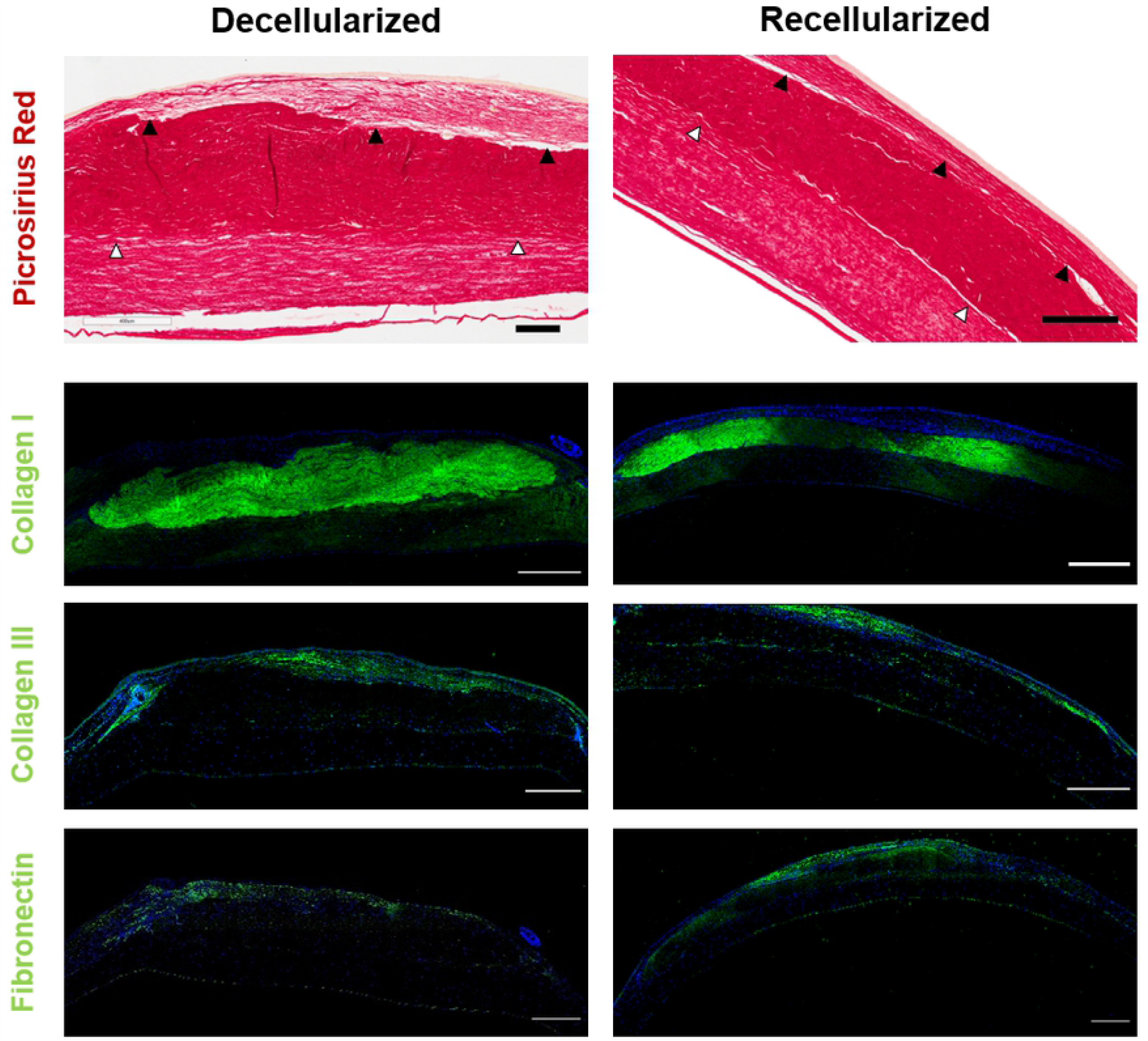
Rabbit cornea 3 months post-implantation stained for cell distribution and phenotype. Representative images of decellularized (left) and recellularized implant (right). A) H&E staining (Scale bar = 1 mm, 200 µm, low and high magnifications). Scaffold apposed with native tissue (white arrows) and regeneration area anterior to the scaffold (black arrows). B) Immunostaining for ALDH1A1 and, C) for α-SMA. B) and C) larger image is whole cornea with close-ups of anterior regenerating layer, central scaffold and posterior native cornea. (Scale bar =500 µm, 50 µm, low and high magnifications).

There was positive cell staining for ALDH1A1 in the epithelium, the native stroma and endothelium (Fig 11B). ALDH1A1 positive cells were also sparsely distributed in the central scaffold area of all of the decellularized and one of the recellularized implants. With α-SMA staining (Fig 11C), there was extensive staining in both groups in the anterior regenerating area, indicating fibrotic cells. Stained cells were also present at blood vessels.

## 4 Discussion

### 4.1 Decellularization effectiveness

There are many decellularization methods that use physical, chemical, or enzymatic methods, often in combination [11]. Decellularization is always a balance; it has to be sufficiently aggressive to remove those cellular components that can cause immunogenic problems following implantation, whilst at the same time being gentle enough to retain the natural environment of the cornea to ensure normal cellularization and function.

Previous studies with the cornea have proposed that a decellularization treatment of NaCl plus a nuclease has advantages over the use of SDS alone. The NaCl method kept the epithelial basement membrane intact and supported the growth of epithelial cells, whereas SDS could not [32]. Also SDS caused high levels of fibril disorganization and poor optical behaviour of these corneas [33]. SDS is effective at decellularization, particularly to remove cell nuclei, but it can be cytotoxic if retained [34]. Combining multiple detergents allows for more complete detergent removal [35] and decreased adverse immune response *in vivo*, but it does increase ECM loss [36], as with the cornea [9] [37]. To promote removal of SDS, Triton-X100 has been used as a second decellularizing agent [35]. In the present study, we used these dual detergents along with dual nucleases to successfully remove over 95% of dsDNA. This achieved levels below the minimal criteria of <50 ng dsDNA mg dry weight, <200 bp DNA fragment length and lack of visible nuclear material in tissue sections stained with DAPI or H&E, that has previously been shown in multiple studies to allow constructive tissue remodelling whilst avoiding adverse cell and host responses [38] [39]. In our study, the decellularization did reduce the GAGs to less than one quarter of that of the native cornea, but the collagen appeared intact. Although SDS does disrupt the basement membrane of human corneas [32], we showed that epithelial cells were able to repopulate the anterior scaffold surface, both *in vitro* and *in vivo*, and our dual decellularization agents and dual nucleases give certainty of adequate cellular immunogens removal. The scaffold also demonstrated adequate optical behaviour with glycerol preparation.

The human cornea accepts allogenic tissue more readily than other tissues, with normal corneal avascularity conferring some immune privilege, such that HLA matching is not usually required if combined with topical immunosuppression. However, humans have pre-formed antibodies to some epitopes, and these can result in the rejection of xenotransplants. For example, native porcine tissue contains galactose-alpha-1,3-galactose (α-Gal) [40] and also N-glycolylneuraminic acid (NeuGc), a non-Gal red blood antigen. These xenoantigens can trigger rejection in pig-to-human implants. In the current study we did not quantify such epitopes, but note that the expression of both Gal and NeuGc have previously been shown to be greatly reduced after decellularization [41]. Additionally, genetically-engineered pigs which express reduced xenoantigens could be used for tissue supply to reduce the human immune response [42].

The detergents used for decellularization can also invoke an immunological problem in their own right. With muscle tissue [43], inadequate extraction of SDS following decellularization resulted in a foreign body response *in vivo*, whereas with a more rigorous extraction the scaffold integrated into the defect with lower levels of inflammatory and fibrosis-related gene expression. In the present corneal study, the scaffold washing method was more intense and of longer duration than the method demonstrated as effective with the muscle, giving in the cornea the same magnitude for the low residual SDS concentration, even before the 28 days of recellularization.

### 4.2 Stromal recellularization

Human keratocytes were used as the cell source for stromal repopulation. The lack of a significant difference between the DNA quantities and the cell depth/density profiles of the three expansion groups appears to indicate that the cell population had reached its peak density within the first 14 days of culture, the short-expansion. Thus, the additional 14 days in the long-expansion or keratocytic-expansion conditions may not be needed to improve cell density, but the extra time may ensure the appropriate phenotype is achieved.

Although after 3 weeks, even with serum [33], keratocytes have been shown to express high levels of ALDH1 protein (a marker of mature keratocytes), unfortunately serum also activates them to become more fibroblastic. To recover quiescence, i.e. an *in vivo*-like phenotype, the serum was removed from the medium in the keratocytic-expansion. This phenotype change was confirmed not only by the expression of the keratocyte cellular markers of ALDH3A1, but also CD34, and the increase of extracellular markers keratocan, decorin and lumican. Under the keratocytic-expansion conditions, the appearance and phenotype of the cells resumed a native form of the uninjured cornea. Immunostaining for α-SMA did not occur in any of the groups, but there was an increase in gene expression with the long-expansion that was even larger with the keratocytic-expansion, indicating the cells were showing signs of moving to a myofibroblastic form.

### 4.3 Clinical outcomes in a rabbit ALK model

Decellularized porcine corneal scaffolds implanted into the human eye have shown different rates of stromal repopulation, ranging from multiple cells within 2 months [2] to little repopulation at 3 months [1]. The depth of the tissue removed and, consequentially, the amount of host stroma remaining may play a role in scaffold repopulation kinetics. These studies demonstrated that keratocyte repopulation can be slow and porcine decellularized tissue has different post-operative healing compared with fresh, cellularized, human cornea [1]. Thus the recellularization used in the present study might be expected to improve healing. Our dissection and positioning of the scaffold maintained its orientation so that the anterior surface was also anterior in the host. Thus the collagen fibre density according to depth should offer a similar biomechanical profile. Nonetheless, in our study with a rabbit host, stromal recellularization prior to implantation did not have much effect on the scaffold’s survival or appearance *in vivo* over 3 months. Both groups appeared to show a regeneration area anterior to the scaffold. The lack of porcine collagen I, but the presence of collagen III in this area indicates new tissue, not reorganised scaffold, as the antibody we used for collagen III reacts against both human and rabbit, but the acellular scaffold had no human cells. It appears from this that the collagen III was from a rabbit cell source.

The phenotype of the cells in the scaffold differed from the cells in the regenerating tissue and the surrounding stroma. The phenotype in the scaffold varied with some cells staining positive for the keratocyte marker ALDH1A1, some for the myofibroblastic marker α-SMA and others for neither marker, potentially indicating a fibroblastic phenotype.

### 4.4 Epithelium restoration but stromal opacity

When a tissue is damaged, a fibrotic response is usually activated. While this response heals, it often fails to restore full function. However, in some instances such as with the cornea, healing can take a regenerative capacity whereby full function is restored, including transparency [17]. Corneal reaction to injury can be attributed to the type of wound. Scarring can be a pathogenic outcome from corneal surgery, and the appropriate epithelial repopulation and basement membrane regeneration can be key to minimize this [44]. It is known that stromal surface irregularities can induce stromal haze and promotion of myofibroblast transformation [45]. In addition the degree of epithelial basement membrane integrity and the extent of epithelial cell cover is an important determinant of whether damage to the stroma is followed by fibrotic or regenerative repair [46]. The epithelial basement membrane acts as a barrier limiting the access of pro-scarring and inflammatory cytokines, such as TGF-β1 and IL-1β, from the epithelium and tears into the stroma [46]. The barrier is achieved by several ECMs, such as nidogens, perlecan or laminin, that are part of the basement membrane of the corneal epithelium and can bind TGF-β1 [47]. Decellularization detergents change the composition of the basement membrane [48], by reducing GAG and other components, thus compromising its barrier function. With tissue replacement, the loss of the epithelial basement membrane has been indicted to allow the free passage of these cytokines into the stroma, where they can activate keratocytes to become myofibroblasts which lay down the scar tissue that results in poor vision. In the present study, scaffolds were covered with epithelium, in agreement with other reports of 3 weeks to confluence [49] [50]. However, this delay in re-establishing the barrier between epithelium and stroma appears to be long enough for the cytokines to trigger adverse change in the keratocytes. In previous studies where the defect was small (2 mm diameter) [2], there was little haze, whereas with a similar treatment but a larger defect (6 mm diameter), the haze was greater [3], as in the present study. This difference may be attributed to a lower cytokine load via the smaller diameter hole. Once the basement membrane is fully regenerated, no more activation of myofibroblasts occurs, and these undergo apoptosis and so, with time, the keratocytes may revert the scar [51]. In the study of Hashimoto et al. 2019 [6], the 8-week corneal appearance was similar to the current study, but at 24 months, corneal transparency was improved. From this we might expect that if our implants had been left in place for longer there may have been optical improvement.

### 4.5 Reinnervation will be dependent upon scaffold organisation

Nerves are severed during human corneal transplantation. Following the use of human tissue allografts, some stromal nerves can appear after 2 months, but basal epithelium re-innervation might not occur until 2 years post-surgery, with corneal sensitivity being recovered no earlier than 3 years [52] [53]. In the present study rat neurites were able to extend though the full depth of the scaffold within 14 days *in vitro* and so the scaffold does not appear cytotoxic, but the environment will differ *in vivo*. In a previous study of decellularized porcine scaffold, nerves only returned in 2 out of 27 cases at 6 months and in only 3 more by 12 months [1]. Nevertheless, it would appear that the extent of reinnervation in porcine scaffold may approach that of native human allografts. Furthermore, if a collagen hydrogel is the implant, then nerve sensitivity can return much more quickly [54]. This may occur because the soft hydrogel permits reduced resistance to penetration and, if this is the case, reinstating keratocytes with a phenotype that can best restore the proteoglycan ground substance between the collagen fibrils may be of benefit.

### 4.6 Alternative methods to address damage from epithelial cytokines

Pocket implants, where the epithelium and its basement membrane are left in place, are not suitable for many surgeries. With ALK, damage to the epithelial basement membrane occurs during decellularized scaffold implantation and this can be detrimental to keratocyte function. Various avenues to prevent this are available which include: 1) re-epithelization *in vitro* to pre-operatively restore the epithelial basement membrane to the implant, 2) prior to implantation, constituent compounds of the epithelial basement membrane, such as laminin, could be layered onto the scaffold to form a smooth barrier to movement of adverse epithelial and tear cytokines before the epithelial layer has time to restore, or 3) drug treatment might be used to counteract these cytokines, allowing keratocytes to migrate from the host, but avoiding their transformation to produce scar tissue. Of these three avenues, prior re-epithelialization with amniotic epithelial cells has been employed [55], but this reintroduces cells and the associated potential immunogenic problem. Adding laminin has also been used in experimental work with compressed collagen gels [56], hyaluronic acid hydrogels [57] and 3D bioprinted corneas [58], but further studies are required. Drug treatment may present thee best opportunity to provide the time for the host epithelial layer to re-establish along with its basement membrane and also time to allow keratocytes to migrate from the host whilst avoiding transformation into myofibroblasts. We have previously shown that biochemical cues can be used to modulate the human corneal stromal cell phenotype to prevent myofibroblast differentiation and in some cases increase proliferation [59]. With corneal endothelium, a TGF-β inhibitor SB431542 has also been used to counteract fibroblastic phenotypes [60], and aid stromal re-innervation [61]. Such techniques of biochemical inhibition and manipulation have potential to augment the use of porcine decellularized corneal scaffold.

## 5 Conclusions

We successfully obtained scaffolds from decellularized porcine corneas with a low DNA and contaminate content that readily supported the growth of epithelial cells and innervation of rat nerve cells *in vitro*. The technique also allowed recellularization of keratocytes deep into the scaffold with a keratocyte-like phenotype and this may also be possible in a shorter timeframe than our 28-day culture. However, with the *in vivo* ALK rabbit model, there was little difference in the ophthalmic outcome between a decellularized and recellularized scaffold in the 3 month time frame of the study. If the study had been over a longer time-course the eventual optical outcome might have improved, but in humans, a period of corneal haze for this length of time, possible scar tissue and an extended period to clarity, is unacceptable for routine use. We conclude that recellularization of decellularized pig cornea with keratocytes may not be sufficiently beneficial to warrant introducing allogeneic stromal cells, unless the stromal environment can also be controlled, including limiting the cytokine influence from the epithelial cells and tears.

## 6 Acknowledgements

The research is supported by funding from the European Research Council (ERC) under the European Union’s Horizon 2020 research and innovation program (grant agreement no. 637460) and from Science Foundation Ireland (15/ERC/3269). We thank the Dublin City University for animal facilities.

## 7 Author Contributions

Conceptualization: MA.

Data curation: JF PWM MA.

Formal analysis: JF PWM.

Funding acquisition: MA.

Investigation: JF PWM RTB PFN.

Methodology: JF PWM MA.

Project administration: MA PWM.

Resources: MA.

Supervision: MA.

Validation: JF.

Visualization: JF.

Writing – original draft: PWM.

Writing – review & editing: PWM JF MA RTB PFN.

